# Tissue- and cellular-level allocation of autotrophic and heterotrophic nutrients in the coral symbiosis – A NanoSIMS study

**DOI:** 10.1101/353250

**Authors:** Thomas Krueger, Julia Bodin, Noa Horwitz, Céline Loussert-Fonta, Adrian Sakr, Stéphane Escrig, Maoz Fine, Anders Meibom

**Affiliations:** laboratory for Biological Geochemistry, School of Architecture, Civil and Environmental Engineering, Ecole Polytechnique Fédérale de Lausanne (EPFL), CH-1015 Lausanne, Switzerland; The Mina and Everard Goodman Faculty of Life Sciences, Bar-Ilan University, Ramat-Gan 52900, Israel; The Interuniversity Institute for Marine Sciences, Eilat 88103, Israel; Electron Microscopy Facility, University of Lausanne, Biophore, CH-1015 Lausanne, Switzerland; Center for Advanced Surface Analysis, Institute of Earth Sciences, University of Lausanne, CH-1015 Lausanne, Switzerland

**Author notes:** Corresponding author: Thomas Krueger.

## Abstract

Corals access inorganic seawater nutrients through their autotrophic endosymbiotic dinoflagellates, but also capture planktonic prey through heterotrophic feeding. Correlating NanoSIMS and TEM imaging, we visualize and quantify the subcellular fate of autotrophic and heterotrophic C and N in the coral *Stylophora pistillata* using stable isotopes. Six scenarios were compared after 6h: autotrophic pulse (^13^C-bicarbonate, ^15^N-nitrate) in either unfed or regularly fed corals, and heterotrophic pulse (^13^C-, ^15^N-labelled brine shrimps) in regularly fed corals; each at ambient and elevated temperature. Host assimilation of photosynthates was similar under fed and unfed conditions, but symbionts assimilated 10% more C in fed corals. Photoautotrophic C was primarily channelled into host lipid bodies, whereas heterotrophic C and N were generally co-allocated to the tissue. Food-derived label was detected in some subcellular structures associated with the remobilisation of host lipid stores. While heterotrophic input generally exceeded autotrophic input, it was more negatively affected by elevated temperature. The reduced input from both feeding modes at elevated temperature was accompanied by a shift in the partitioning of C and N, benefiting epidermis and symbiont. This study provides a unique view on the nutrient partitioning in corals and highlights the tight connection of nutrient fluxes in symbiotic partners.

## Introduction

The immense biodiversity and productivity of tropical coral reefs exists in a marine environment that is generally characterized by low nutrient levels and plankton concentrations. Coined as “Darwin’s paradox”, the investigation into why corals can thrive in areas with high wave action and low nutrients has directed the scientific focus towards nutrient fluxes and cycles within the overall reef framework and within individual hermatypic corals (1, 2). The coral animal has access to the inorganic nutrient pool in the surrounding seawater through endosymbiotic photosynthesizing microalgae (mainly *Symbiodinium* sp.) and via parts of its microbial community, such as cyanobacteria and diazotrophic bacteria (3, 4). Intracellular *Symbiodinium* sp. cells photosynthesize in the light and fix CO_2_ into triose phosphate compounds that can serve as C skeletons for subsequent processes, such as inorganic N assimilation (5). The CO_2_-fixation rate in *Symbiodinium* is very high and C is rapidly assimilated into sugars and amino acids within seconds to minutes (6, 7). The translocation of part of these soluble photosynthates from the symbiont to the coral tissue happens across the entire colony surface, depending on local symbiont density. While it has been demonstrated that coral autotrophy can be sufficient to meet the metabolic C demands of the coral animal (8, 9), coral heterotrophy (e.g. hunting for prey) is also an important component of coral nutrition, especially in cryptic and mesophotic habitats or turbid water environments, where light becomes a limiting factor (10-12). Upon loss of autotrophic input, as is the case in bleached corals, heterotrophy becomes fundamental for survival and recovery of the animal (13, 14). Under normal, healthy conditions, the relative contribution of autotrophy and heterotrophy to coral nutrition is modulated by the environment and may vary between species (15). Importantly, the intracellular microalgae, in addition to assimilating inorganic nutrients from the seawater, also allow for a reassimilation of metabolic ‘waste’ products of the animal host (e.g. NH_4_^+^ and CO_2_) (16). Hence, the symbiotic nature of the coral animal ensures both access to organic and inorganic nutrients and recycling and conservation of acquired nutrients in an oligotrophic environment.

Despite the autotrophic capability of scleractinian corals, which originated in the middle to late Triassic (17-19), reef-building symbiotic corals have retained highly developed feeding mechanisms (behavioural and anatomical features) that characterize them as heterotrophic organisms. Coral recruits have been observed to capture zooplankton as early as 8 days after settlement (20). Three forms of heterotrophic feeding have been categorized by Goreau, Goreau (21): (i) “specialized carnivorous feeding, primarily on zooplankton”, (ii) “unspecialized detritus feeding”, and (iii) “direct utilization of dissolved and colloidal organic matter”. Coral polyps actively prey on zooplankton using cnidocyte containing tentacles and/or mesenterial filaments (22). Ciliary currents directed towards the mouth or the generation and re-ingestion of mucus nets and strings on the surface can furthermore enhance prey capture (23, 24). All coral polyps in colonial corals are connected by coenenchyme tissue (*sensu* coenosarc tissue) and their gastrovascular cavities are linked by a network of gastrovascular canals that serve as a circulatory system for transporting material across the colony (25-27). Despite their wide-ranging gastrovascular systems, corals lack specialized organs for storing molecules that fuel metabolism. Instead the coral tissue contains a high density of transitory C storage bodies, such as lipid bodies and small granular glycogen deposits that all receive photoautotrophic C (7, 28, 29).

The assimilation of nutrients generally serves two purposes: to generate energy (catabolism) or to build cellular structures (anabolism). Excretion of metabolic waste products and secretion of products of anabolism (mucus production, reproductive output) constitute the main pathways for the loss of nutrients in corals (28, 30). Quantifying and modelling the partitioning of nutrients from autotrophic and heterotrophic pathways between host and symbiont has traditionally been achieved through measurements of radioactive or stable isotopes in bulk tissue samples (3, 31-34). More recently, by mapping the subcellular distribution of elements and their stable isotopes, nanoscale secondary ion mass spectrometry (NanoSIMS) has opened a new visual dimension to coral nutrition and has captured the dynamic nature of nutrient fluxes within and between different coral tissue compartments (7, 35–39). In contrast to bulk measurements, NanoSIMS analysis provides not only ultrastructural information on the fate of nutrients when correlated with transmission electron microscopy (TEM) imaging, it also provides a unique anabolic view on coral metabolism and the relative turnover of elements within specific structures.

Here, we used NanoSIMS to contrast the relative turnover and identify the primary sinks of autotrophic and heterotrophic nutrients in the surface body wall of the coral coenenchyme in the scleractinian coral *Stylophora pistillata.* We tested the extent to which the compartment-specific assimilation rates from both feeding modes are modulated by elevated temperature and whether a regular food supply to the coral host alters the autotrophic importance of its symbionts with regard to C and N assimilation and translocation.

## Material and Methods

### Coral maintenance and experimental design

Specimens of *Stylophora pistillata* were collected from the shallow waters (4-8m) in the Gulf of Aqaba in Eilat, Israel under permit 2013/40158 of the Israel Nature and Parks Authority. Experiments were conducted in the Red Sea Simulator Red Sea aquarium setup at the Interuniversity Institute (IUI) for Marine Sciences and corals maintained in outdoor flow-through aquaria (30L; >60L h^−1^) under ambient water temperature and reduced solar irradiance (ca. 350 μmol m^−2^ sec^−1^ at midday). Small aquaria pumps close to the water surface provided additional flow and turnover in each aquarium, while breaking the surface to increase light scattering. Three different mother colonies were fragmented to provide six similarly sized clonal fragments per colony. Fragments were subsequently acclimated to the aquarium settings for 3 weeks and then each colony allocated to four treatments (ambient *vs.* elevated temperature with or without feeding) in a paired design and maintained over 67 days (Fig. S1). The heat-treated fragments (originating from colonies A, G, I) underwent a thermal profile with a cumulative heat exposure of 11.2 degree heating weeks (DHW) at ambient pH 8.1, identical to the experiments of Krueger, Horwitz (39), which took place in parallel. Corals in the feeding treatments were fed twice a week with freshly hatched *Artemia* nauplii (2500 per coral fragment) for one hour with interrupted seawater flow. The daily mean temperature in the ambient and elevated treatment for the ten days leading up to the experimental incubation was 24.1±0.2°C and 29.1±0.2°C, respectively (39).

### Isotopic labelling of seawater for autotrophic pulse

Seawater for the autotrophic labelling pulse was spiked with 2 mM NaH^13^CO_3_ (98 atom %) and 3 μM K^15^NO_3_ (98 atom %). The chosen 3 μM nitrate spike for the 6 h labelling period brings the total nitrate concentration to approximately ten times higher than natural values for this location (0.2-0.3 μM) and most other reef locations (40, 41). It was used here to spike and track ^15^N within a 6 h period, but nevertheless represents a nutrient level that does naturally occur in some reef locations, e.g. near equatorial upwelling regions (e.g. Galapagos islands; 3-5 μM), in nearshore regions of the Great Barrier Reef and Coral Triangle (~1-3 μM), and in some reefs in the Gulf region (e.g. Strait of Hormuz, Gulf of Oman; 2-3 μM) (41–43)

### Isotopic labelling of Artemia sp. for heterotrophic pulse

*Tetraselmis* sp. cultures that would later serve as food source to label *Artemia* sp. were started in modified f/2-medium (-Si), using 0.22 μm-filtered natural seawater that was supplemented with 2 mM NaH^13^CO_3_ and in which the nitrate source was replaced with Na^15^NO_3_ (0.882 mM final concentration). Cultures were grown in this medium for 3 weeks, with occasional sub-culturing. Final ^13^C- and ^15^N- enrichment of a lyophilized aliquot were factors of 105 and 182 times higher than the natural ^13^C/^12^C and ^15^N/^14^N ratios, respectively. *Artemia* cultures were fed daily with this labelled *Tetraselmis* culture for a total period of nine days after hatching. The feeding period of *Artemia* with labelled algae was limited to nine days in order to obtain considerable enrichment with labelled C and N, while avoiding that *Artemia* individuals became too large to serve as life prey for *Stylophora pistillata. Artemia* cultures were maintained in a 30 L outdoor aquarium tank that was also supplemented with 2 mM NaH^13^CO_3_ and 3 μM Na^15^NO_3_ to minimize loss of labelling in remaining *Tetraselmis* cells and label other phytoplankton that potentially grew in this feeding tank. At the end of the labelling period all *Artemia* individuals were collected with a plankton net and concentrated in 2.5 L to assess the size of the population and determine the feeding density for the experiment. A bulk *Artemia* subsample that underwent the same resin embedment as the actual coral samples was measured in the NanoSIMS and the overall ^13^C- and ^15^N-enrichments were 3.65 and 138 times the natural isotope ratios, respectively. The purpose of this was to relate the measured coral enrichment to the initial food enrichment (see section Data representation and normalisation), thus also accounting for dilution effects of the resin on the isotopic enrichments of these samples.

### Experimental incubations

Incubations with labelled seawater or labelled food were done in the same 30 L aquaria that were used for the acclimation period. However, in and outflow was interrupted for the labelling period and water movement solely provided by the heat exchanger and the surface pump. To assess whether the autotrophic assimilation is affected by regular host feeding, one set of corals that had been acclimated to regular feeding in both temperatures was moved into the aquaria used for the autotrophic labelling pulse one day before the experiment. Thus, three different incubation modes were performed: autotrophic pulse with unfed corals (Aut [U]), autotrophic pulse with regularly fed corals (Aut [F]), and heterotrophic pulse with regularly fed corals (Het [F]) (Fig. S1). Coral fragments in both temperature treatments were exposed to labelled seawater or labelled living *Artemia* (~1059 individuals per fragment) for a total of 6 hours (10 am to 4 pm local time) with a replacement of water every two hours to minimize the increase in water temperature due to the high ambient air temperature. Density of labelled *Artemia* was monitored by counting individuals from 30 mL water samples in a zooplankton counting chamber (Hydrobios Apparatebau GmbH, Altenholz, Germany) under a binocular every 20 mins. After adding life *Artemia* prey to the aquarium, corals developed mucus threads and many *Artemia* got visibly trapped or directly captured by coral polyps; there were no individuals in the water column after 80 mins.

### Sample preparation, TEM, and NanoSIMS

At the end of the isotopic pulse, the apical tip of the coral branch was removed and a ca. 1 cm coral piece clipped off and immersed in fixative (2.5% [v/v] glutaraldehyde, 0.5% [v/v] formaldehyde in 0.22 μm filtered 0.1 M phosphate buffer with 0.6 M sucrose, pH 7.4-7.6) for 2 h at room temperature, followed by 22 h at 4°C. A piece of a unlabelled coral colony from fed and unfed conditions was used as reference to assess the natural isotopic ratio in the tissue of interest. Pieces were subsequently washed and completely decalcified in 0.5 M EDTA. Fixed tissue samples were dissected to obtain connective coenenchyme tissue and post-fixed in 1% [v/v] osmium tetroxide in water for 1 h. We exclusively analysed the coenenchyme tissue between polyps in order to avoid data artefacts due to the individual polyps catching different amounts of prey within the pulse period. All used histological terminology follows Peters (22). Post-fixed samples were dehydrated, embedded in Spur resin blocks, microtomed and post-stained for TEM, following established protocols (7). Transmission electron microscopy (TEM) was carried out at 80 or 100 kV with a Philips CM 100 and a Tecnai 12 (FEI) transmission electron microscope, respectively. While ultrathin sections (70 nm) were mounted on carbon formvar-coated Finder copper grids and imaged for correlative TEM and NanoSIMS, semi-thin sections (500 nm) were mounted on glass slides and directly imaged with NanoSIMS in order to cover larger areas and to acquire sufficient data for statistical analysis.

Suitable areas were analysed for ^13^C- and ^15^N-enrichment using a NanoSIMS 50L ion microprobe. Sections were gold-coated and bombarded with a 16 keV primary Cs^+^ ion beam. Raster scans of the area of interests were performed with a beam focus spot size of around 150 nm (6 layers). Image resolution was set to 256×256 pixels with a beam dwell time of 5 ms per pixel. Image size for semi thin sections was set to 40×40 μm, which provided a complete cross section of the surface body wall epidermis and gastrodermis with sufficient resolution for subcellular features to be clearly resolved. Image sizes for correlative TEM and NanoSIMS varied. For semi-thin sections, approximately 10-15 images were obtained per replicate in order to cover enough area to obtain enrichment values for 40-60 symbiont cells and their surrounding host tissue. In case multiple sections from the same sample block were required to obtain enough cells, sections were taken at least 20 μm apart to avoid cutting and imaging another layer of the same symbiont cells. The secondary ions ^12^C_2_^−^ (mass 24), ^13^C^12^C^−^ (mass 25), ^12^C^14^N^−^ (mass 26), and ^12^C^15^N^−^ (mass 27) where simultaneously counted in electron multipliers at a mass resolution of about 10000, enough to resolve all potential interferences.

### Data treatment and normalisation

NanoSIMS data were processed using the L’IMAGE software (created by Dr. Larry Nittler, Carnegie Institution of Washington). Maps of ^13^C/^12^C and ^15^N/^14^N ratio distributions were derived from drift corrected image ratios of ^13^C^12^C^−^ and ^12^C_2_^−^, and ^12^C^15^N^−^ and ^12^C^14^N^−^, respectively. TEM and corresponding NanoSIMS images were used to visually identify enriched subcellular structures. Note that shown TEM images were adjusted for contrast. In order to permit easy comparison between different NanoSIMS images, the enrichment levels in ^13^C and ^15^N for all presented images are expressed using the δ-notation, which expresses the relative difference between the measured isotope ratio in a labelled tissue (R_sample_) and the measured natural isotope ratio in the same tissue type (R_reference_) in parts per thousand:

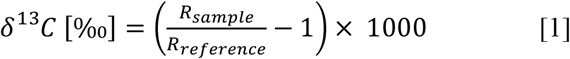

Regions of interest (ROIs) were drawn on ratio images to assess average enrichment in each type of compartment. The following ROIs were defined and manually drawn on each image based on contour outlines visible on the ^12^C^14^N^−^ image and/or the isotopic ratio images: Symbionts (cross-sectional diameter >3 μm), extra-algal lipid bodies (outside of symbionts, but contained within the symbiosome), host gastrodermis (excluding symbionts), host epidermis, and host lipid bodies (diameter >1 μm). Each NanoSIMS image thus provided data points for multiple symbiont and lipid ROIs and one data point for gastrodermis and epidermis, respectively. All NanoSIMS images were slightly smoothed (smooth width of 3 pixels).

In order to be able to accurately contrast the input from autotrophic and heterotrophic nutrients, the measured enrichment in each ROI was normalized to the isotopic labelelling of the respective source (seawater or brine shrimp) as outlined in the following. For subsequent calculations, the measured isotopic ratios were converted into ^13^C and ^15^N atom fractions, *F* (in atom percent, AP, when expressed as *F*× 100):

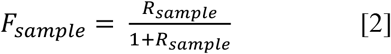

The atom ratios for the resin embedded heterotrophy food source were determined with *F*_Het_C_ = 0. 04772±0.01155 and *F*_Het_N_ = 0.33772±0.01360 (N=6). Atom fractions of ^13^C and ^15^N in the spiked seawater (*F*_SW_C_ and *F*_SW_N_) were calculated taking into account the relative concentrations and atom
fractions of nitrate and C in seawater and the added spike. Assuming standard atom fractions for C (*F*_pdb_: 0.01111) and N (*F*_N_: 0.00366) (44) in the natural seawater and ambient concentrations of 2.2 mM total dissolved inorganic C and 0.3 μM nitrate (40), the following equations express the final seawater atom fractions in 1 L of seawater after the addition of the spike (*F*_SW_). Atom fractions for the added spike were 0.98 for both NaH^13^CO_3_ and K^15^NO_3_, as indicated by the supplier (Sigma-Aldrich, St Louis, MO, USA).

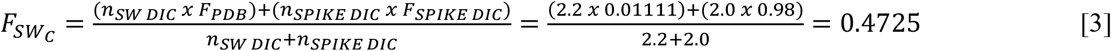

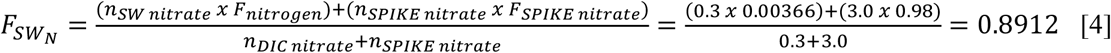

Using these *F*_SW_ values, the relative atom percent excess (APE) in autotrophic samples was calculated for each element as:

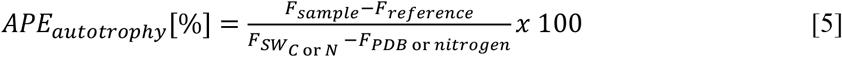

Using the measured atom fraction from labelled and unlabelled *Artemia* in resin, the corrected APE for the heterotrophic coral samples was calculated for each element as:

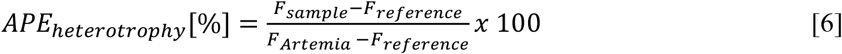

In essence, these APE for each type of ROI express the proportion of the total C or N pool that has been replaced with newly assimilated C and N from either food source. Assuming steady state conditions of the C and N pool, i.e. assuming a negligible change in total biomass in the imaged coenenchyme areas within the pulse period, APE is a measure of the relative C and N turnover in each ROI over 6 h in the light. In general, it should be noted that conventional preparation of biological samples for NanoSIMS involves a loss of most soluble compounds and restricts observations mainly to the anabolic part of metabolism. Thus, the presented values should be interpreted as minimal estimates for tissue C and N assimilation.

Since APE only represent the relative turnover of a compartment without considering the actual size of this compartments, a semi-quantitative measure of C and N assimilation was calculated by multiplying size of compartment (cross-sectional area A, in μm^2^) with relative turnover in this compartment (APE in %) for each element. This “integrated area fraction” (IAF) for each type of compartment allows an assessment of relative C and N partitioning across the main three tissue compartments (sym: symbiont, gastr: gastrodermis, epi: epidermis) of the surface body wall within the 6 h pulse, where the total assimilated C or N pool is composed of:

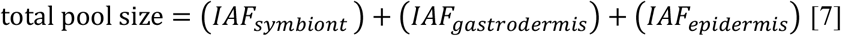

Note, that IAF is a semi-quantitative measure based on the reasonable assumption that the C/N elemental ratio in these compartments is similar, consistent with NanoSIMS images (data not shown). For a fully quantitative comparison, one would have to determine the total C and N-pool in each compartment separately. Note that IAFs and the resulting partitioning budget only considers the structurally incorporated C and N; i.e. it does not consider losses due to mucus shedding or respiration.

### Statistical analysis

Considering that ROI enrichment values contain two sources of variability, resulting from the cutting level within the cell (affecting for example the abundance of starch granules in a particular symbiont ROI) and the true biological variability as result of size, age and physiological state of individual cells, we decided to treat the ROI data points as raw data points and include biological replicate as a factor in the analysis, rather than treating multiple ROIs from one biological replicate as technical replicates and reducing them to a single value. Thus, we sought to balance the number of data points for the same types of ROI between the three biological replicates for statistical considerations. Note that the published unfed autotrophy dataset from Krueger, Horwitz (39) was reused here and contrasted with other feeding conditions. Depending on the specific question, the three datasets (Aut [U], Aut [F], Het [F]) were tested individually or jointly with multifactorial ANOVAs for each ROI type. In case of insignificant interaction terms, the model was reduced to a minimal adequate model. Significant interaction terms were analysed with Tukey HSD *post hoc* tests. How regular host feeding affects the autotrophic assimilation at different temperatures in each compartment was tested with a Three-Way ANOVA with temperature, feeding acclimation, and replicate as factors (Aut [U] vs. Aut [F]; Table S1). Secondly, the heterotrophy dataset (Het [F]) was tested for the effect of temperature (Table S2). In the third analysis, we tested whether there is a difference in the tissue-specific C and N contribution from autotrophy and heterotrophy and how elevated temperature interacts with this. This was tested with a Three-way ANOVA for each compartment, with temperature, feeding mode and replicate as factors (Aut [F] *vs.* Het [F]; Table S3). In addition to the species’ mean response, individual responses of the colonies are displayed since there were usually significant differences in the C and N-turnover between replicates (raw data summary in Table S5).

## Results

### Ultrastructural observations

Autotrophically fixed C in the symbionts was mainly concentrated in the primary starch ring around the pyrenoid, in secondary starch granules, in lipid bodies throughout the symbiont cell, and as translocated product in the lipid bodies of the host gastrodermis (Fig. 1A, B, D, E). In addition, occasionally detected extra-algal lipid droplets, located outside of the algae, but within the symbiosome (Fig. 1A, B: LB*), showed very high levels of C-turnover (8-11%). Fixed nitrate was incorporated throughout the symbiont cells at high levels, outlining them against the host gastrodermis (Fig. 1C, F). Occasionally, strongly labelled ^15^N hotspots were observed inside these cells. Within the symbionts, the accumulation body was the structure with the lowest levels of photosynthetic C or N incorporation.

The host tissue showed different degrees of labelling with autotrophic N and the highest enrichment was usually observed in the immediate vicinity of symbiont cells, where host nuclei and nucleoli showed significant N incorporation (Fig. 1C). The symbiosome membrane complex that engulfs the *Symbiodinium* cells also incorporated a large amount of fixed nitrate (Fig. 1C, F arrow).

**Figure 1.**
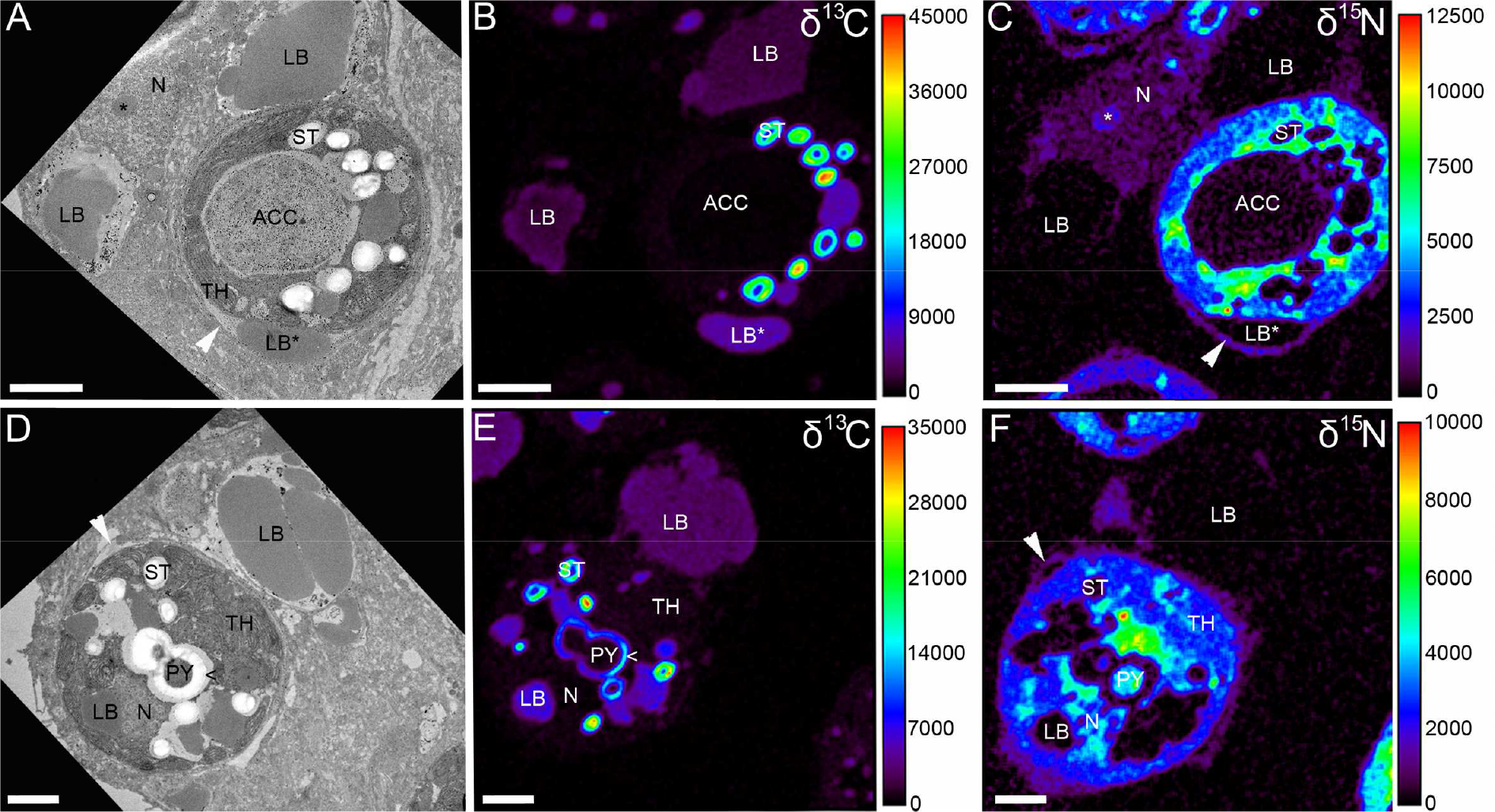
Autotrophic uptake of carbon and nitrogen. Correlated TEM and NanoSIMS images of the distribution of ^13^C and ^15^N in *Symbiodinium* and surrounding gastrodermal cell cytoplasm, resulting from simultaneous photosynthetic fixation of [^13^C]bicarbonate (B, E) and assimilation of [^15^N]nitrate (C, F) after a 6 h isotopic pulse in the light. Fixed ^13^C and ^15^N was detected throughout the symbiont cell, with carbon mainly concentrated in the primary starch sheet (<) around the dinoflagellate pyrenoid (PY) near thylakoid membranes (TH), in secondary starch granules (ST), symbiont and host lipid bodies (LB), and in extra-algal lipid bodies (LB*). Symbiont nitrogen labelling was lowest in carbon-rich structures and the accumulation body (ACC), and highest in small crystalline-like hotspots. Translocated ^13^C to the host tissue was mainly concentrated in lipid bodies, whereas nuclei (N) and nucleolei (*) in the host cell showed high ^15^N labelling. Colours in NanoSIMS maps represent enrichment relative to an unlabelled tissue in δ-notation (black to red). White arrows (A, D, C, F) point to symbiosome membrane. Scale bars are 2 μm.

After 6 h of heterotrophic feeding, ^13^C and ^15^N derived from the digested brine shrimp was detected in all coral tissue layers, including the gastrodermis of the basal body wall and the calicodermis (Fig. 2, S2A-C). Enrichment was highest in the gastrodermis for both elements, where numerous gastrodermal hotspots (~1-2 μm diameter) expressed strong co-labelling of C and N at both temperatures (ambient/elevated: Spearman’s ρ=0.75/0.81, both p < 0.0001, N=490/173). Three distinct types of labelled hotspots could be differentiated in the gastrodermis: (i) round, membrane-delineated vesicles that contained homogenous labelled material (pink arrows, Fig. 2, 3D, 4), (ii) elongated, clearly defined electron-dense structures, that were of cup shape and in direct contact with the lipid bodies (blue arrows, Fig. 2, 3E, 4, 5), and (iii) oval or elongated patches, which were difficult to distinguish from the surrounding tissue apart from being electron-lucent (green arrows, Fig. 2, 3F, 4). With regard to their heterotrophic isotopic signature, vesicles and electron-lucent patches formed a continuum, whereas the cup-shaped structures tended to be more heavily enriched in ^15^N relative to ^13^C (Fig. 4D-F).

**Figure 2.**
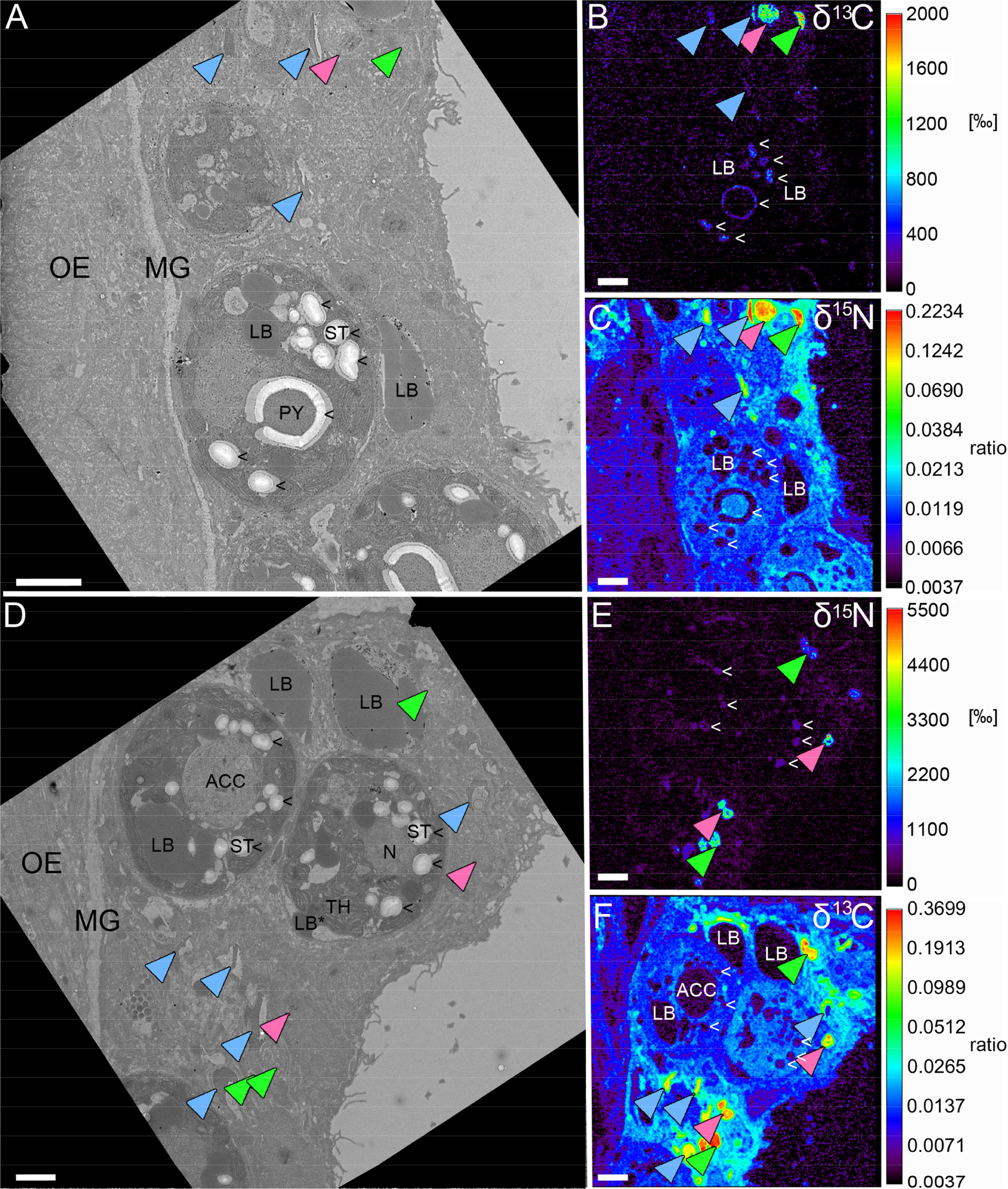
Heterotrophic carbon and nitrogen assimilation of coral gastrodermis. Correlated TEM and NanoSIMS images of the distribution of prey ^13^C (A, E) and ^15^N (C, F) in the surface body wall gastrodermis 6 h after ingestion of labelled *Artemia salina* began. Subcellular features abbreviated as in Fig. 1 with OE indicating the surface body wall epidermis and MG the mesoglea. Multiple gastrodermal ^13^C and ^15^N hotspots were observed (coloured arrows, more detailed in Figs. 3, 4). Hotspots of ^13^C in the symbionts corresponded to starch (<). Colours in NanoSIMS maps represent enrichment relative to an unlabelled tissue in δ-notation (B, E, black to red) or as ratio (C, F; note log scale). Scale bars are 2 μm.

**Figure 3.**
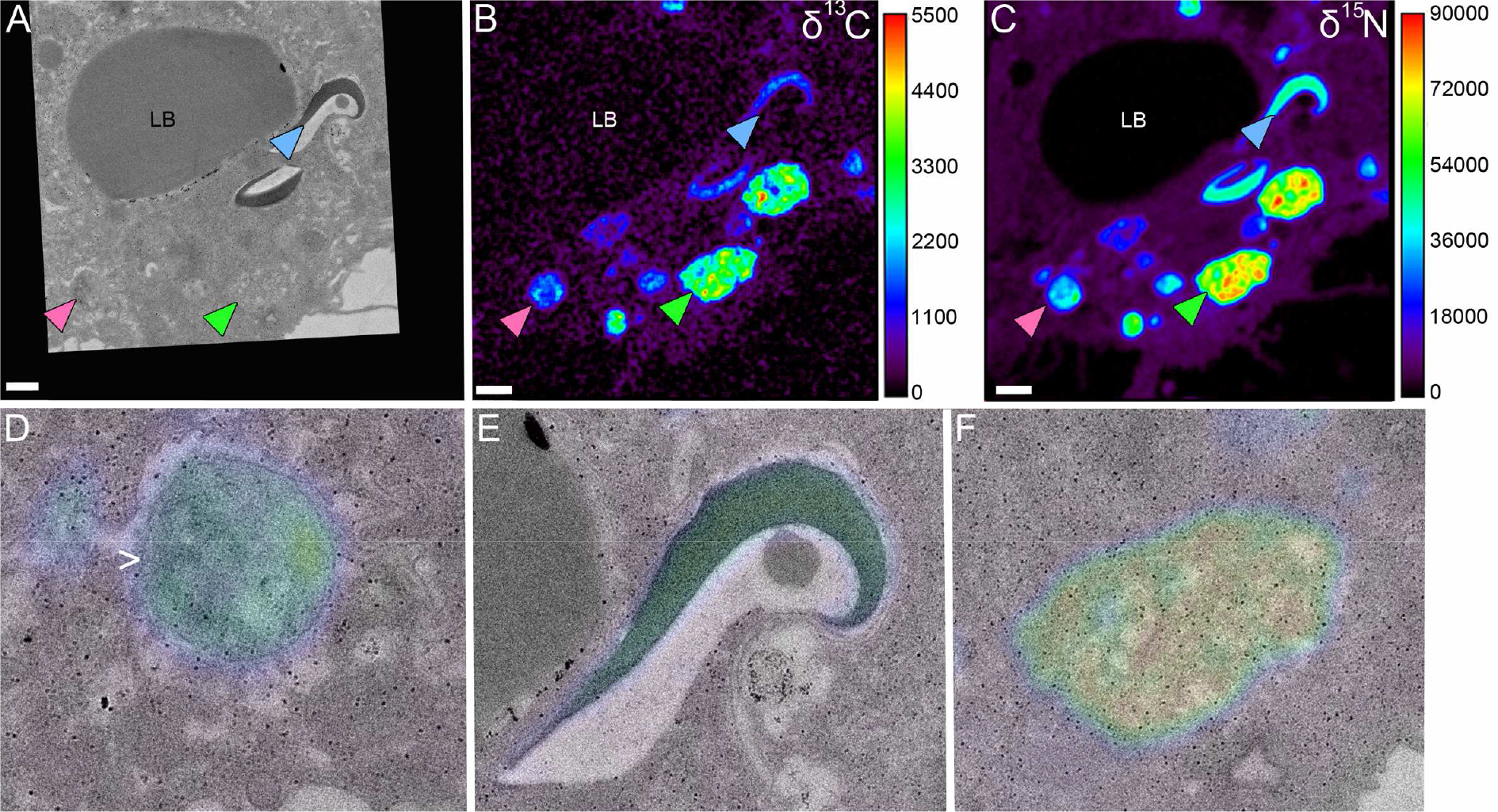
Types of isotopic hotspots in the gastrodermis from heterotrophy. ^13^C (B) and ^15^N (C) enrichment maps of gastrodermis exhibiting three different types of isotopic hotspots (indicated by colours; see text for discussion). (D-F) TEM ultrastructure and δ^15^N NanoSIMS overlay. Note the presence of a membrane around the hotspot in D (>). Colours in (B) and (C) display enrichment relative to an unlabelled tissue in 5-notation (black to red). Scale bars are 1 μm.

**Figure 4.**
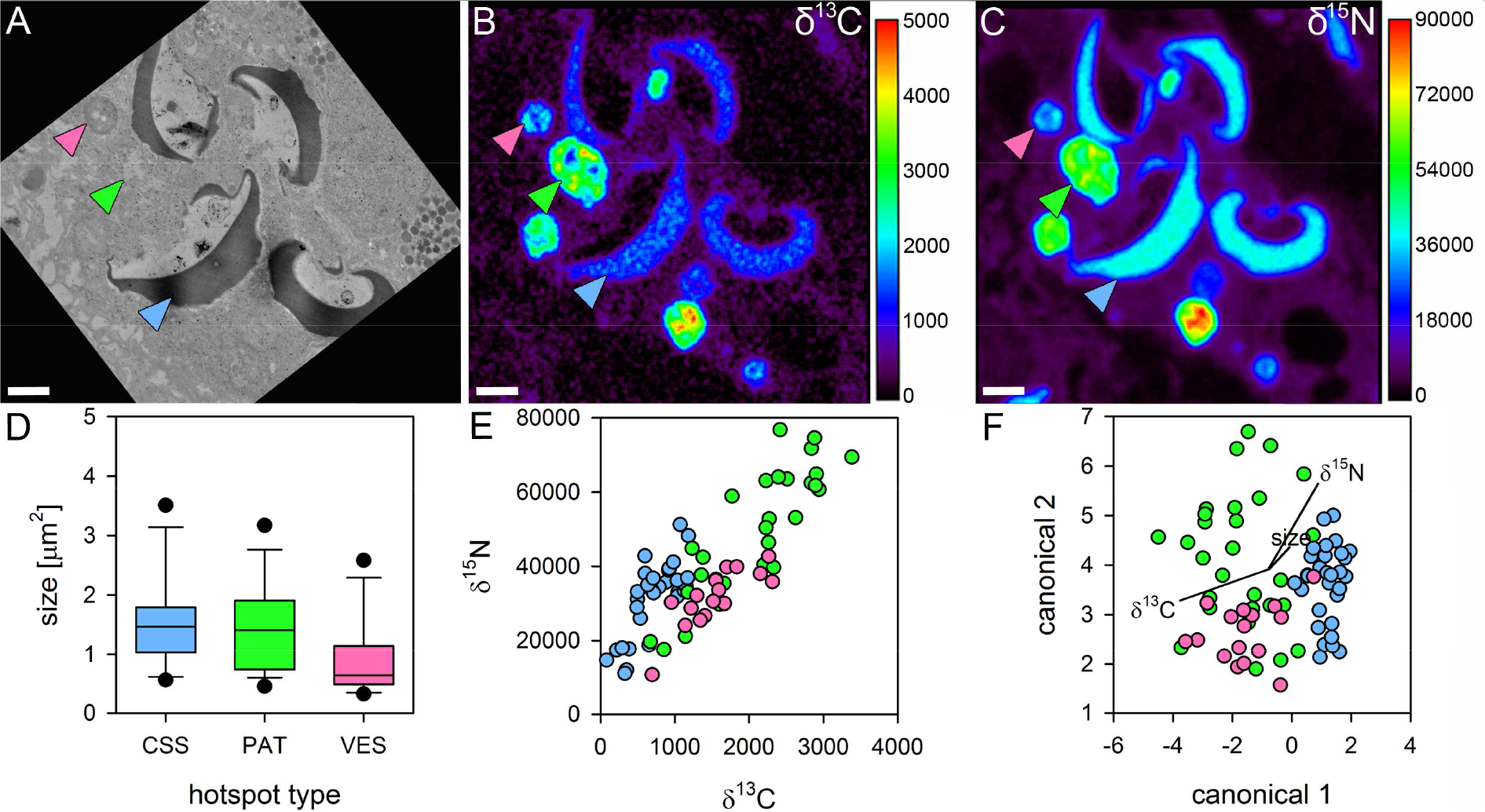
Characteristics of isotopic hotspots in the gastrodermis from heterotrophy. Multiple high-resolution (15×15 μm) TEM and NanoSIMS images (A-C) were analysed for size (D) and individual enrichment (E) of the three types of isotopic hotspots in the gastrodermis. CSS cup-shaped structures (blue arrows), PAT electron-lucent patches (green), VES vesicles (pink). Canonical correlation analysis of these characteristics revealed a distinct CSS cluster with overlapping PAT and VES clusters (F). Colours in (B) and (C) display enrichment relative to an unlabelled tissue in 5-notation (black to red). Scale bars are 1 μm.

**Figure 5.**
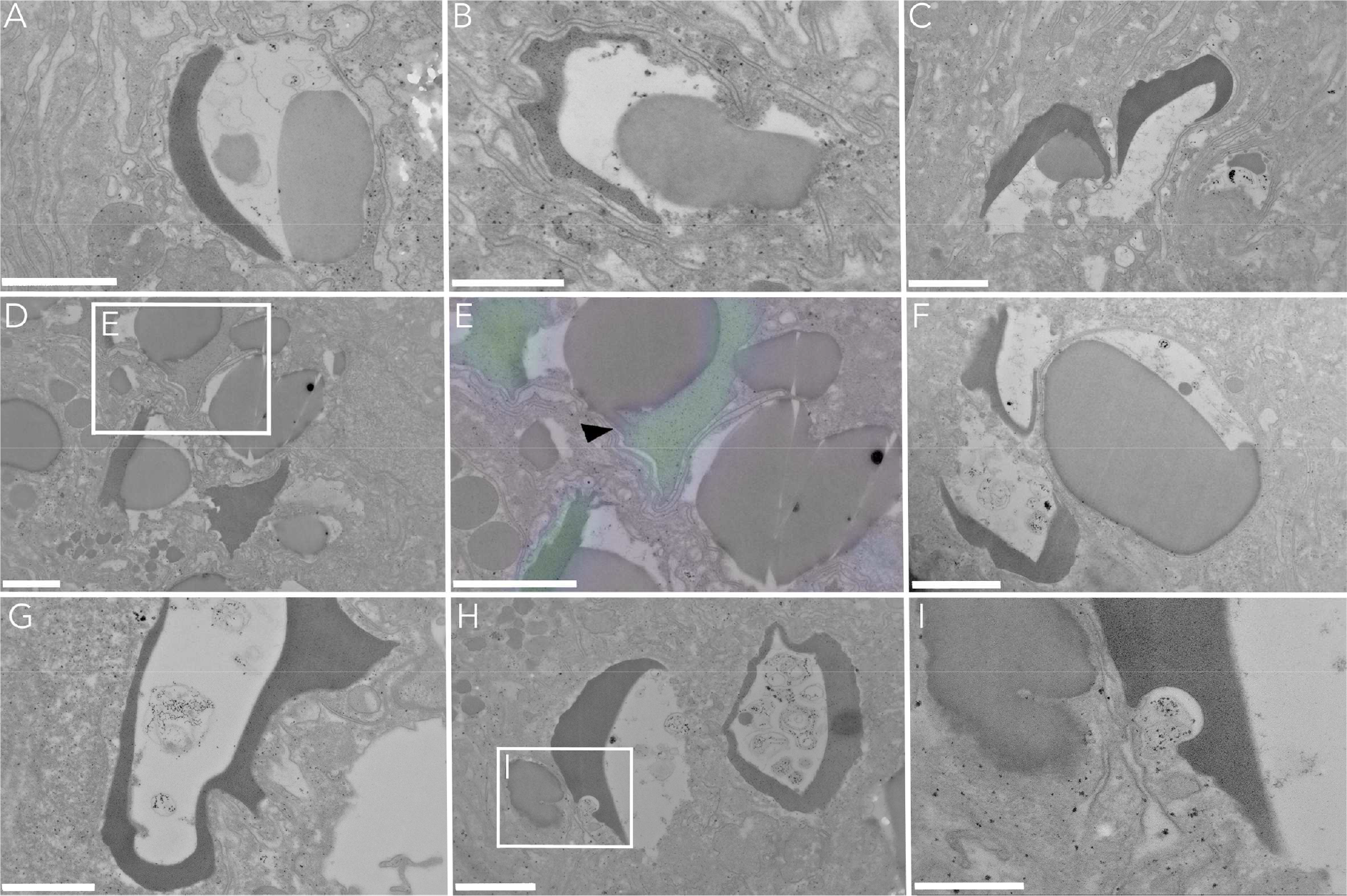
Association of cup-shaped structures with gastrodermal host lipid bodies. Cup-shaped electron-dense structures that were highly enriched with heterotrophic C and N (cf. Fig. 4) were consistently found next to partially extracted host lipid bodies (A, B) or in direct contact with them (C-E). E shows the formation of a membrane continuity between both structures, despite the distinct separation of the isotopic signal (black arrow in E with δ^15^N-signal as colouration). Areas with extracted lipids consistently showed the presence of residual material (F-H). Intrusion of the host ER membrane into the cupshaped structure (I). Scale bars are 1 μm (A, C-H) or 500 nm (B, I).

A few isotopic hotspots were also found in the host epidermis, representing mostly small vesicles, but with a much lower level of labelling than the gastrodermal vesicles (Fig. S2D-F). The association of cup-shaped structures with host lipid bodies was very prevalent (Fig. 5A-E) and usually accompanied with varying degrees of lipid degradation, creating empty spaces that contained residual material (Fig. 5 F-I). In one case we even observed the formation of a membrane continuity between both organelles, despite the clear spatial separation of the isotopic enrichment (Fig. 5D, E). Heterotrophic C and N were also detected in the symbiont after 6 h. Here, N-labelling throughout the symbiont cell was homogenous, although occasional small vesicle-like hotspots were observed (Fig. S3). Detectable C labelling was limited to starch granules and the primary starch sheet around the pyrenoid (Fig. 2A, B, D, E).

Average symbiont cell size *in hospite* across the three replicates was 7.3±0.1 μm (mean±SE, total N=316) at ambient temperature and they occupied 30-40% (N=160) of the gastrodermal crosssectional area. A significant increase in average cell size was detected at elevated temperature, further increasing when combined with feeding (temperature x feeding acclimation: *F*_1,605_=4.21, p=0.0407*). Average symbiont cell diameter in fed corals at elevated temperature was with 8.1±0.2 μm (N=153) significantly larger than under all other conditions (based on *post hoc* Tukey HSD).

### The effects of regular feeding and temperature on coral autotrophy

The symbiont and the host epidermis were the only structures for which regular feeding showed an effect on the turnover of autotrophic nutrients, specifically C. Symbionts in corals that experienced regular feeding showed significantly increased autotrophic C acquisition by 10±8% at both temperatures (mean±SD, N=6, Fig. 6A, Table S1). Such a positive effect was also seen for epidermal C turnover, albeit much smaller and only at ambient temperatures (+51±57%; N=3; Fig. 6C, Table S1). In general, elevated temperature significantly lowered the turnover of autotrophic C and N in all main coral compartments for both elements, except for N in the host lipid bodies (Table S1, Fig. 6). Structural incorporation of autotrophic C and N at elevated temperature declined on average by 19±12% and 5±26% in the symbiont, irrespective of feeding state. Likewise, gastrodermal C and N incorporation declined significantly (−40±29% and −29±36%), with epidermal N turnover dropping on average by −33±16%. Autotrophic C turnover in the host tissue was highest in the host lipid bodies with ~8% C replacement over 6 h in the light, while at the same time displaying a similar N turnover as the overall gastrodermis (Fig. 6D, F, H). For host lipid bodies, elevated temperature only significantly affected C (−12±17%, N=6), but not N turnover.

**Figure 6.**
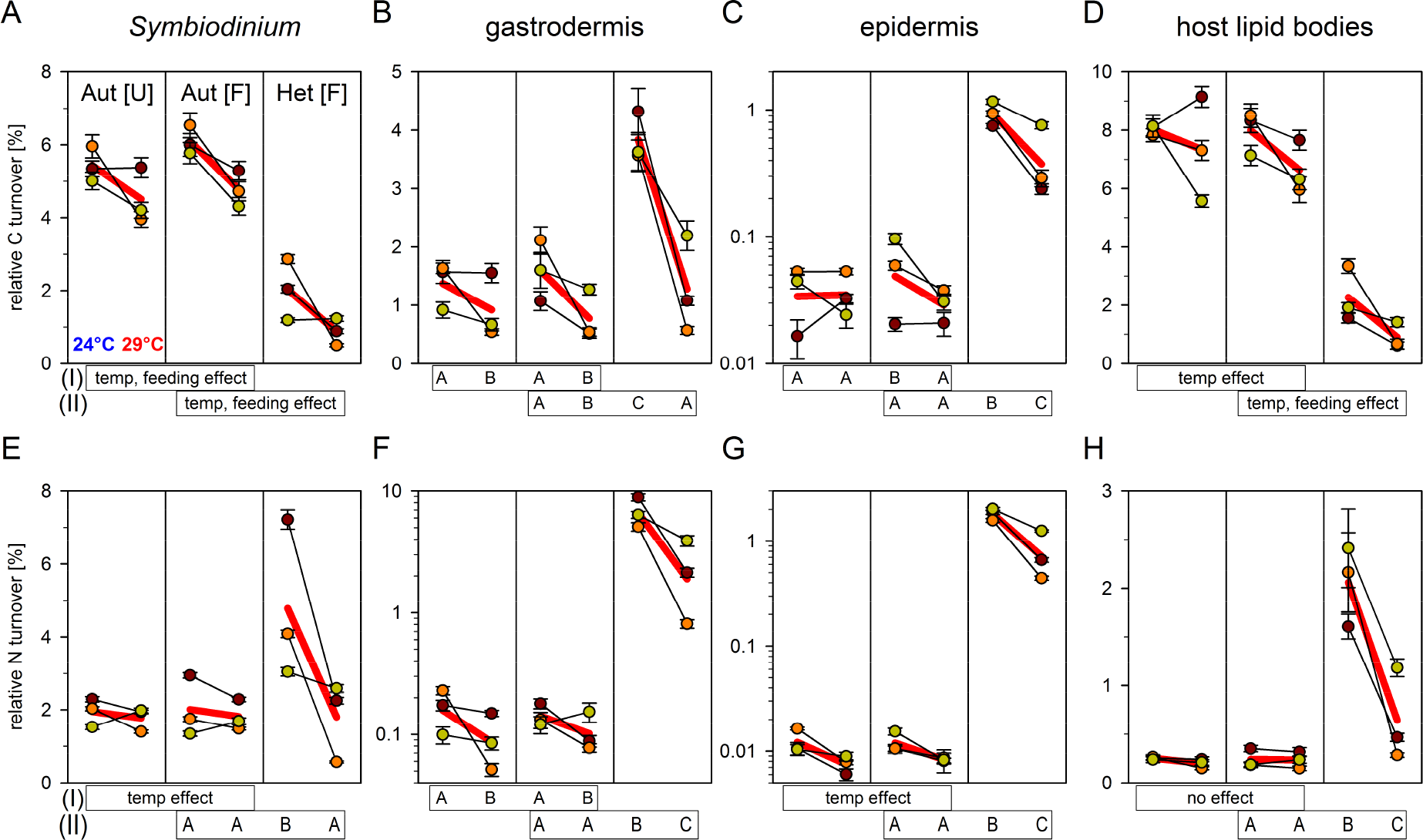
Compartment-specific carbon and nitrogen turnover. Shown is the paired response of three Northern Red Sea *Stylophora pistillata* coral colonies at the end of a 6-week exposure period to ambient (24°C; data to the left in each column) or elevated SST (29°C; data to the right in each column). Temperatures indicate average values for the last 10 days leading up to the measurements. Individual colony responses (dots; green, orange, red) and overall species response (thick red lines) are shown for *Symbiodinium* and different host structures of the surface body wall of the coral coenenchyme. Aut [U] indicates autotrophic (Aut) assimilation of bicarbonate and nitrate by corals unfed ([U]) for 67 days. Aut [F] and Het [F] indicate autotrophic and heterotrophic (Het) assimilation, respectively, in corals fed ([F]) regularly twice a week prior to the measurements. Relative C and N turnover values (see Material & Methods) represent the proportion of the total C or N pool that has been replaced with newly assimilated ^13^C and ^15^N from each food source, assuming steady state conditions. These data show the degree of C and N-assimilation in each compartment, independent of its absolute size. Note log scales in C, F, G. Data points are averages obtained from 40-60 imaged symbionts and their surrounding tissue (mean±SE; Table S4). Statistical effects of (I) temperature and feeding acclimation in Aut[U] vs. Aut [F] (cf. Table S1) and of (II) temperature and feeding mode in Aut [F] vs. Het [F] (cf. Table S3) on C and N turnover are shown below each subpanel. In case of significant interactions of the two main factors, Tukey HSD *post hoc* results are depicted as capital letters, where treatments not connected by the same letter are significantly different.

Partitioning of autotrophic C and N between symbionts and the two host tissue compartments of the surface body wall was very similar between regularly fed and unfed corals, and temperature had no substantial effect on this (Fig. 7). After 6 h, approximately 75% and 90% of the detected C and N was located in the symbiont tissue that made up ca. 1/5^th^ of the cross-sectional area of the surface body wall. The remaining material was mainly located in the gastrodermis and only 1-2% of the autotrophic C and N pool was present in the epidermis, despite representing half of the area of the surface body wall coenenchyme. The disproportional retainment of C and N in the symbionts meant also that allocation of C to the gastrodermal layer was at least twice as high as N allocation (~25% vs. ~10%, Fig. 7).

### Autotrophic vs. heterotrophic C andNanabolism under ambient and elevated temperature

After 6 h of photosynthetic activity, the symbionts replaced on average 5-6% of their structural C (from DIC) and ~2% of their N (from nitrate) under ambient conditions (Fig. 6A, E). Average structural C and N-turnover in the gastrodermis due to translocated photosynthates was with 1.0-1.6% and 0.1-0.2%, respectively, approximately ten times higher than in the host epidermal layer, which only showed little acquisition of photoautotrophic nutrients (Fig. 6B, C, F, G).

Heterotrophic nutrient input considerably exceeded autotrophic input for the gastro and epidermal compartment, especially for N, where turnover-values where one to two orders of magnitudes higher under ambient temperatures (Fig. 6, Table S3). Heterotrophic N contribution to the symbiont population after 6 h also exceeded its own nitrate fixation by a factor of two to three, while C incorporation was approximately three times lower than the autotrophic input (Fig. 6A, E). In contrast to the autotrophic feeding mode, heterotrophic C was not concentrated in host lipid bodies, which showed lower C enrichment than the overall gastrodermis and received four times less C from heterotrophy compared to autotrophy.

Elevated temperature had a distinct effect on the C and N-metabolism of the coral compartments. Temperature consistently reduced autotrophic and heterotrophic input to all compartments (Fig. 6, Table 1, S1, S2), and these temperature-induced drops in nutrient assimilation were significantly larger for the heterotrophic feeding mode (with the exception of C-assimilation in symbionts and host lipid bodies; Fig. 6, Table 1, S3). This strong impact of temperature led to a situation where heterotrophic assimilation rates dropped by 50-70% and even fell to the level of autotrophic rates observed at ambient temperature in two of the compartments (C in gastrodermis and N in symbiont; Fig. 6B, E). Concomitantly, significantly reduced C− and N− enrichment under elevated temperature was also observed in the heterotrophic hotspots (Fig. S4).

**Table 1.** The effect of temperature on autotrophic and heterotrophic input. Shown are the mean changes in autotrophic (Aut) and heterotrophic (Het) carbon and nitrogen turnover for each compartment (mean±SD, N=3) in regularly fed [F] corals (cf. Fig. 6). All compartments refer to the surface body wall of the coral coenenchyme. Pairs with asterisk indicate that temperature caused a significant stronger reduction in heterotrophic than autotrophic turnover (Fig. 6, Table S3).

|  | Aut [F] vs. Het [F] |  |
| --- | --- | --- |
|  | <sup>13</sup>C | <sup>15</sup>N |
| *Symbiodinium* | -22±9% vs. -45±44% | -4±26% vs. -57±37%* |
| host gastrodermis | -49±27% vs. -66±24%* | -22±43% vs. -66±24%* |
| host epidermis | -34±35% vs. -57±19%* | -29±15% vs. -58±18%* |
| host lipid bodies | -17±12% vs. -55±27% | 0±25% vs. -70±18%* |

Comparison of the autotrophic and heterotrophic nutrient partitioning between the three tissue compartments of the surface body wall after 6 h of incubation showed radically different pictures for both feeding modes. Heterotrophy caused a strong co-allocation of C and N with almost identical patterns of C and N partitioning across the three compartments (Fig. 7). While the majority of the autotrophic C and N pool is retained in the symbiont after 6 h (see above), approximately two thirds of the heterotrophic nutrient pool is present in the gastrodermis after 6 h under ambient conditions. The remaining third is approximately equally split between symbiont and epidermal compartment, matching the relative compartment size in the case of the symbiont (Fig. 7). The epidermis is the main benefactor of the heterotrophic feeding mode and received a substantially larger share for both elements under ambient conditions compared to autotrophy (~21% *vs.* ~1%; Fig. 7). The effect of elevated temperature was not just restricted to a reduction in overall nutrient assimilation, but also caused a considerable shift in the partitioning of nutrients, especially for the heterotrophic feeding mode. Here, the shift primarily benefited the symbiont and epidermal compartment and reduced the share of the gastrodermis from originally ~63% to ~50%. This pattern of increasing allocation towards 20 symbiont and epidermis to the disadvantage of the gastrodermis was also observed in the autotrophic dataset of regularly fed corals, albeit much more subtle (Fig. 7).

**Figure 7.**
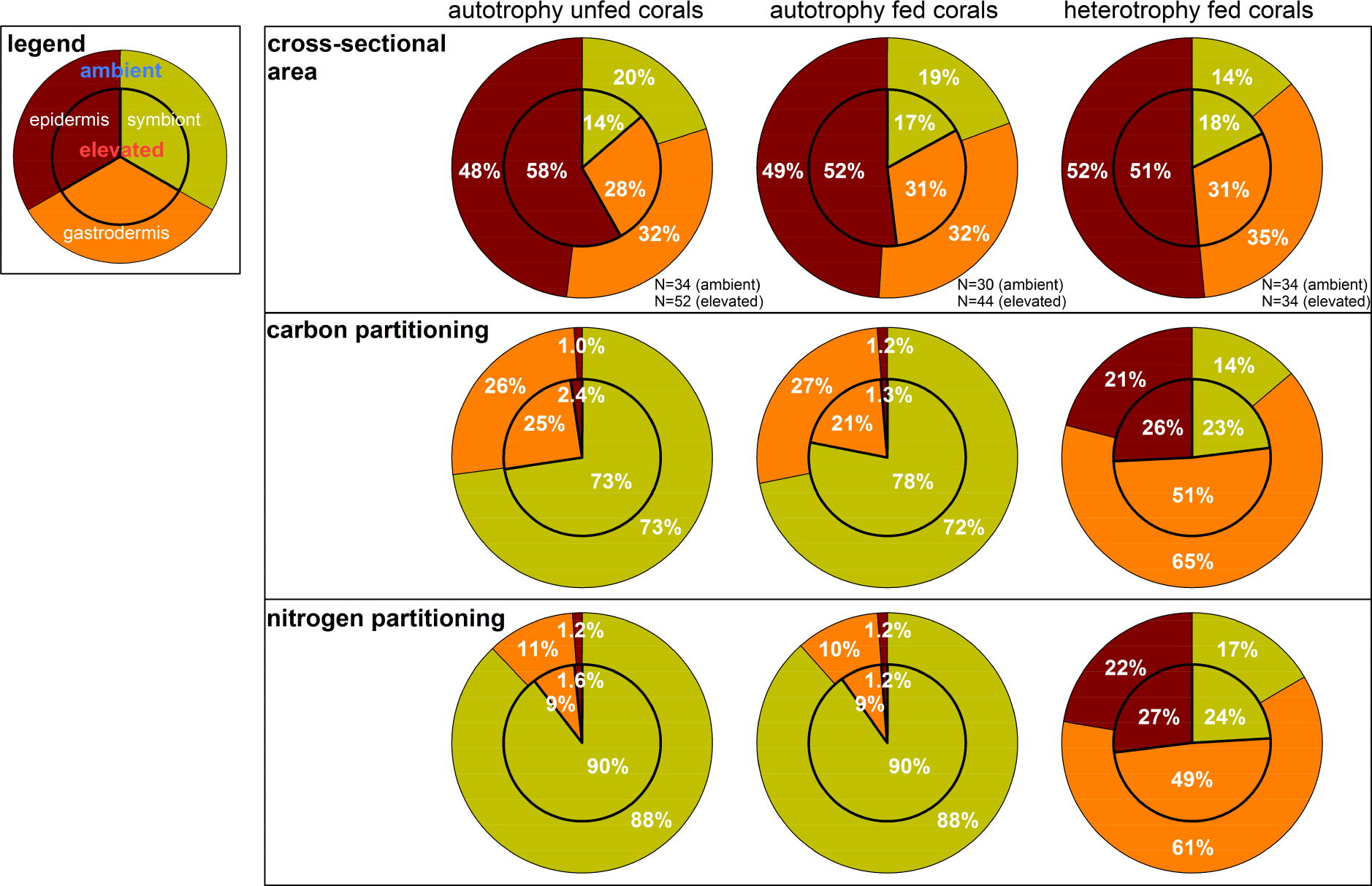
Partitioning of assimilated ^13^C and ^15^N in the coral symbiosis after 6 hours. Shown are the relative fraction of the cross-sectional area of each compartment of the surface body wall coenenchyme and the ^13^C− and ^15^N distribution across each compartment in each feeding mode (pie charts left to right) and at both temperatures (inner and outer pie chart). Data for C and N partitioning were obtained by combining the cross-sectional area of each compartment with its ^13^C and ^15^N turnover (Fig. 6) obtained from multiple NanoSIMS images (see IAF calculation in Material & Methods). These data provide a semi quantitative view on the distribution of the assimilated C- and N-pool at the end of the 6 h pulse.

## Discussion

By employing correlative TEM-NanoSIMS isotopic imaging, our study revealed the primary sites for the concentration and storage of nutrients derived from auto and heterotrophy and highlights the critical role that feeding mode and seawater temperature have on the relative turnover and partitioning of C and N within the different tissues of the intact symbiosis. The following discussion will almost exclusively make reference to prior studies of scleractinian corals. Although sea anemones are considered good coral ‘model organisms’ for many research questions, they are not suitable models for nutritional research of the coral symbiosis. Anemones are characterized by a thick macroscopic tissue and absence of a skeleton, creating different demands and utilization dynamics for C and N compared to a coral animal with a thin tissue layer over a massive C skeleton sink. In the same way, assimilatory performance of the symbionts, e.g. with regard to nitrogen, differs in anemones due the fundamentally thicker host tissue (e.g. absence of detectable nitrate fixation [(45), personal observation] due to high levels of metabolic ammonia release). Thus, we will primarily consider previous studies of auto and heterotrophy from scleractinian corals for comparison and discussion.

### The ultrastructural fate of autotrophic nutrients

Consistent with previous studies, the correlative TEM-NanoSIMS imaging revealed that fixed DIC was deposited primarily in starch granules and lipid bodies, whereas nitrate was distributed relatively homogenously throughout the symbiont cells of *Stylophora pistillata* (35, 37). The occasional ^15^N hotspots were ultrastructurally difficult to interpret, but likely resembled the previously characterized uric acid crystal-containing vesicles in *Pocillipora damicornis*, which serve as highly concentrated transitory N stores (37). Nevertheless, these ^15^N-hotspots were present at relatively low density and absent from some symbiont cell sections, likely due to the lower concentration of the nitrate spike used here (3 μM instead of 30 μM). The repeated observation of extra-algal lipid bodies (i.e. contained within the symbiosomal vacuole, but outside the algal cell) with a high level of ^13^C-enrichment (cf. Fig. 1A, B this study, 7, 46, 47) confirms the release of photosynthetically derived lipids into the host vacuole. ^13^C and ^15^N from translocated photosynthates were detected in all coral layers, but large gastrodermal lipid bodies served as the main C stores for the coral host. The ten times higher N enrichment in the symbiont compared to host gastrodermis at the end of the pulse period perfectly matched previous bulk measurements of *Stylophora pistillata* under similar conditions (61); even though corals here fixed twice as much nitrate overall. Distribution of autotrophic ^15^N in the host tissue was relatively homogenous and the general protein turnover in the host tissue. Differential labelling of subcellular organelles was only distinguishable in the immediate vicinity of the symbionts, where translocated ^15^N was most concentrated. Here, stronger ^15^N labelling of host nuclei and nucleoli was clearly visible (cf. Fig. 1C) and especially host nucleoli showed a high turnover for autotrophic and heterotrophic N, likely due to the faster turnover of nucleolar ribosomal proteins relative to the nuclear structure (48). Interestingly, the symbiosome membrane that encapsulates the symbionts in their host-derived vacuole appeared to incorporate a substantial amount of photosynthetically derived N within the pulse period. Indeed, in the small stretched membrane part highlighted in figure 1C, the N turnover was with 0.58% ca. 3-6 times higher than the overall gastrodermal value (cf. Fig. 6F). This rapid turnover of its proteinaceous components reveals a highly dynamic nature of the symbiosome membrane that is fuelled by the photosynthate delivery of its occupant.

### Heterotrophic C and N in the host gastrodermis

Nutrients derived from the digestion of the isotopically enriched brine shrimps in the polyps were detectable throughout the wider coenenchyme tissue. Their particular enrichment in the gastrodermis of the surface and basal body wall that directly face the gastrovascular cavity was consistent with the role of the gastrovascular canals as a distribution system throughout the colony (25). The similar ^13^C- and ^15^N-enrichment values from random coenenchyme patches of three independent coral colonies (Fig. 6) demonstrated a highly efficient distribution and assimilation of heterotrophic food over the entire colony surface after just 6 h. Given that the provided food was of proteinaceous nature, the high degree of co-labelling of C and N in the overall gastrodermis and in the different hotspots as well as the similar partitioning of both elements across the compartments suggests that basic monomers containing both elements (e.g. amino acids) were used directly in anabolic processes, rather than being de-aminated or extensively processed involving internal reserves prior to incorporation into the tissue. Nevertheless, the three observed types of hotspots reflected different stages of food processing. The round membrane-delineated ^15^N-hotspots (pink arrows Fig. 2-4) constitute vesicles that contained food material that was endo or phagocytosed from the gastrovascular canals and transported through the host tissue. These structures are likely of transitory nature and a direct result of the ingestion and processing of heterotrophic food. The second type of ^15^N-hotspot structure, which appeared cupshaped (blue arrows Fig. 2-5), seemed to be more permanent, because it regularly occurred in *S. pistillata* samples and was also observed in adults and larvae of *P. damicornis* (49, 50). Their electron dense appearance in the TEM indicates either a higher physical density or a high affinity for the heavy metals used in the post-fixation and staining, suggesting the presence of a high degree of negatively charged polar groups. Their close association with lipid bodies and the presence of empty space surrounding these structures might indicate a role in the remobilization of host lipid bodies. Their direct contact with host lipid bodies and their rapid incorporation of heterotrophic N and external ammonia suggest a crucial function in the coral lipid metabolism, potentially related to breakdown of lipid droplets as evidenced by the residual material in the vicinity of the extraction. The third type of hotspot (green arrows in Fig. 2-4) usually showed the highest levels of heterotrophic ^13^C and ^15^N enrichment (with ca. 40-60% of the brine shrimp ^13^C and ^15^N enrichment level), but its specific function remains unclear. The high degree of heterotrophic C- and N-enrichment either indicates the lowest degree of degradation and processing of the original food or cellular patches with the fastest turnover that rapidly and extremely efficiently incorporate nutrients into cellular material. Since no clear ultrastructure was distinguishable and such highly enriched gastrodermal hotspots were not found in the autotrophic treatment, we discount however the possibility of dedicated cytoplasmic areas with permanently high C and N turnover. The fact that all these hotspots showed reduced enrichment with labelled brine shrimp material at elevated temperature suggests that all these bodies are involved in the breakdown and storage of heterotrophic nutrients and experienced a lower influx of heterotrophic food.

An interesting observation was the strong gradient between host gastrodermis and epidermis with regard to C and N assimilation in both feeding modes. It appears that the thin collagenous mesoglea acted as a border that restricted flow between both tissue layers and limited the distribution and assimilation of nutrients primarily to the side that faces the gastrovascular canal (including shuttling of food vesicles throughout the gastrodermis). The consistent large difference in C and N turnover between both layers indicated either a significant delay in the anabolic utilisation of nutrients originating from the gastrodermis or a fundamentally different anabolic metabolism, with epidermal cells having a much slower turnover. Notably, we detected substantial enrichment from heterotrophic food also in the basal body wall gastrodermis and the adjacent calicodermis, illustrating a direct contribution of heterotrophically derived nutrients to the calicodermal activity that forms the skeleton.

### Assimilation of heterotrophic nutrients by Symbiodinium

The labelling of primary and secondary starch deposits in the symbiont with heterotrophic ^13^C demonstrated that coral catabolism released brine shrimp C through decarboxylation reactions and the tricarboxylic acid cycle (TCA), which was then directly re-assimilated as CO_2_ in symbiont photosynthesis. Indeed, comparing the fate of labelled C from incubation with 1 mM [1-^13^C]-pyruvate (label released as CO_2_ in the pyruvate dehydrogenase complex) or [3-^13^C]-pyruvate (label enters TCA cycle) over 3 h in a separate experiment, confirmed that the initial decarboxylation of pyruvate prior to entering the TCA cycle is a major source for symbiont C recycling (Fig. S5B). In contrast, fewer ^13^C was photoassimilated from the CO_2_-release of the TCA cycle and the homogenous host tissue labelling from this form indicated that the label entered primarily pathways of amino acid and fatty acid synthesis via cataplerotic reactions of the TCA cycle and direct Acetyl-CoA use, respectively (Fig S5E). The different labelling patterns from these two incubations confirm that catabolic CO_2_ is fixed by the symbiont and that phototrophic carbon translocation is primarily directed towards the host lipid bodies. To what degree host lipid bodies receive phototrophic carbon directly through the use of translocated lipids or indirectly through sugar breakdown and *de novo* fatty acid synthesis via glycolytically derived Acetyl and Malonyl-CoA remains however to be answered.

Analogously to heterotrophic C assimilation by the symbiont, a substantial enrichment of the symbiont with heterotrophic N and the occurrence of occasional N hotspots illustrated a rapid assimilation of brine shrimp N. Our data indicated that ~14-24 % of the assimilated heterotrophic C and N pool was found in *Symbiodinium* after 6 h (based on cross-sectional areal budgets, Fig. 7), roughly matching earlier observations in *Oculina arbuscula* and *O. diffusa* (10-20% of ^15^N over 4-28 h (32, 51)). Since the measurements here were *in hospite*, previous concerns about incomplete homogenization and host tissue contamination of the symbiont pellet as experimental artefacts were avoided. An interesting aspect with regard to the transfer of nutrients from host to symbiont is the question of how fast the symbiont population can access heterotrophic N. Piniak and Lipschultz (51) argued that the occurrence of heterotrophic N in the symbiont within 4 h was insufficient time for a complete recycling of ^15^N from host metabolism (“i.e. host digestion, synthesis into host macromolecules, catabolism, excretion, uptake by zooxanthellae”) and that the symbionts must have assimilated prey N directly. Our data seem to support this notion, considering that the relative amount of replaced N in the symbiont population under ambient temperatures in two of the three colonies was only 19% and 20% lower than the corresponding host gastrodermal values after 6 h (the third one was 52% lower). Considering their intracellular location and the consistent observation of assimilation of organic substrates (and amino acids in particular) by *Symbiodinium* (52-55), these symbionts likely directly assimilate amino acid monomers as well as released ammonium as they appear in the tissue in the process of the breakdown of the brine shrimp material (assuming transport across the symbiosome membrane). Indeed, the NanoSIMS images from auto and heterotrophically ^15^N-enriched symbiont cells were qualitatively similar, notwithstanding the difference in absolute level of labelling (compare Figs. 1C, F and S3B). Thus, in our interpretation, the small ^15^N hotspots in the symbionts represent the immediate concentration and storage of nitrate and ammonium ions into crystalline uric acid (this study, 37), whereas the more or less homogenous labelling of all subcellular symbiont structures represents the incorporation of more complex N-containing compounds from the brine shrimp into the general cellular infrastructure. Despite the additional influx of prey N, internal N recycling by the symbionts is a key mechanism for the coral and previous studies have estimated that up to ~80-90 % of the symbiont N is derived from this process (16, 33). Here, heterotrophic N considerably supplemented the symbiont’s N status (exceeding its own nitrate fixation by a factor of 2.4; Fig. 6A) and this supply in combination with the influx of other limiting elements such as phosphorus likely lifted the general nutrient limitation (56-58).

### Modulation of symbiont autotrophy by holobiont feeding

The motivation to investigate symbiont autotrophy efficiency under different feeding regimes came from the observation that regularly fed corals tend to display higher host tissue thickness and increased symbiont densities that alter the light microenvironment for individual cells (reviewed in 58). While such tendencies were also observed here (Table S4), the NanoSIMS data showed that the symbionts in fed corals assimilated on average 10% more C, but not more N (Fig. 6A, B). Assuming that the relative activity of host anabolism and catabolism was not fundamentally different between unfed and regularly fed corals, this extra amount of photosynthetically assimilated C was primarily utilized by the symbionts themselves similar to the observation in another feeding experiment (12). These observations thus confirm that the enhanced N supply from regular feeding increased individual symbiont C investment into cellular structure (as reflected by the NanoSIMS data and the tendency of increased symbiont size and density in fed corals). Thus, the uncoupling of symbiont photosynthesis and growth under nutrient limitation is eased under regular host feeding, leading to the expected effect that the symbionts retain a larger share of their fixed C for growth (reviewed in 59). We attribute the unchanged state in nitrate fixation to the isometric scaling of increased host tissue biomass and symbiont biomass (symbiont density per host biomass was not significantly affected; Table S4). Thus, the tendency for increased host tissue biomass and elevated ammonia production that might have shifted the N prevalence of the *in hospite* symbiont from nitrate to the preferred ammonia (60–65), as is the case in anemones (45), was counterbalanced by increases in symbiont numbers (p=0.053, Table S4). For the effect of elevated temperature, we can conclude that the feeding state of the holobiont made no difference for the generally reduced autotrophic acquisition of C or N.

### Coral autotrophy vs. heterotrophy and the impact of elevated temperature

Our data clearly demonstrated that the influx of heterotrophic C and N to host tissues substantially exceeded any photosynthate contribution, in particular for N from nitrate fixation (Fig. 8). When extrapolated from these numbers, the single feeding event immediately (within 6 h) provided the host gastrodermis with the C and N equivalent of ~14 and ~283 daylight hours worth of translocated photosynthates. This gradient was even stronger for the epidermis, with the C and N received from the single feeding event equating to ~97 and ~903 daylight hours of photosynthetic input, respectively. Although an earlier model calculation estimated that approximately only one third of the daily N demand is covered through nitrate assimilation in *Stylophora pistillata*, heterotrophic N influx would still considerably exceed the symbiont’s inorganic N fixation (NO_3_^−^ and NH_4_^+^ make up 75% of the daily budget) (66). Thus, while symbiont C is mainly derived from seawater DIC, internal recycling and heterotrophic input are the main source for symbiont N (Fig. 8). Our NanoSIMS data demonstrated that whereas photoautotrophic C is mainly directed towards storage in the host lipids,heterotrophic C is primarily assimilated into the host tissue (Fig. 8). We suggest that this was likely due to particular proteinaceous nature of the ingested zooplankton, with a limited deamination of amino acids to generate keto acids and ketone bodies that would be used in glycolytic or TCA-linked pathways to form Acetyl-CoA for lipogenesis.

**Figure 8.**
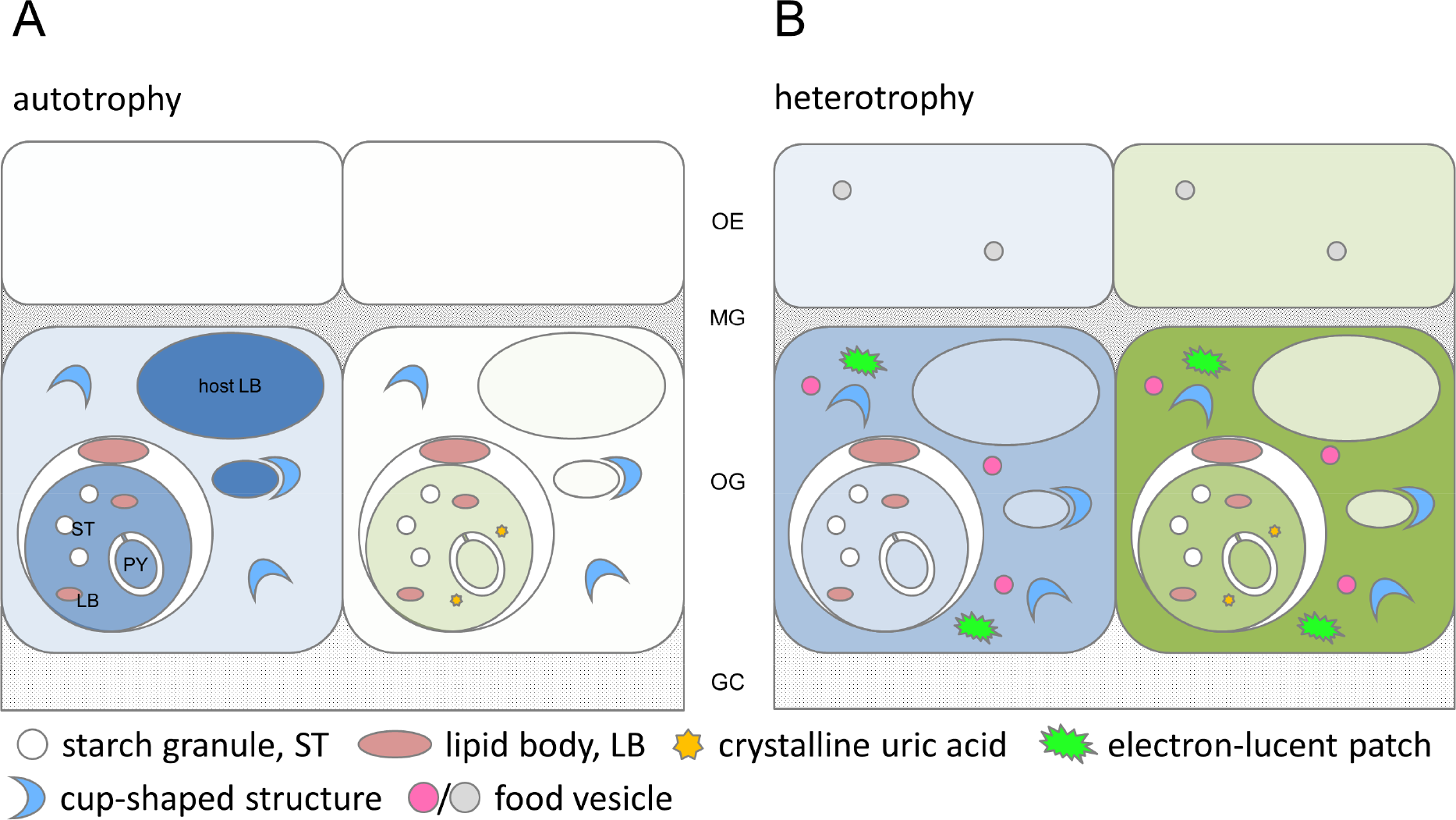
Schematic overview of nutrient input and ultrastructural features involved in the autotrophic and heterotrophic feeding mode of *Stylophora pistillata*. Influx of carbon (left cell; blue) and nitrogen (right cell; green) into symbiont, host lipid bodies, gastrodermal, and epidermal cell, originating from symbiont photoautotrophy (A; seawater bicarbonate and nitrate) or host heterotrophy (B; zooplanktonic prey) after 6 h. Colour tone contrasts the relative influx from both feeding modes for C (blue) and N (green), respectively. Subcellular structures involved in storage and processing of nutrients are highlighted. OE, oral epidermis; MG, mesoglea; OG, oral gastrodermis; GC, gastrovascular canal.

As previously reported, negative effects of elevated temperature (11.2 DHW) in this Red Sea *Stylophora pistillata* were largely restricted to lowered autotrophic nutrient turnover and not reflected in most other physiological variables (39). This observation also applies to the heterotrophic feeding mode. Although still providing a net influx of nutrients, turnover on the tissue-level scale showed even stronger temperature-induced reductions for N (all compartments) and C assimilation (both host tissue layers) from the heterotrophic feeding mode compared to the autotrophic mode. The simplest explanation for the reduced assimilation could be a reduced feeding rate at elevated temperature as a previous study on a Red Sea *Stylophora pistillata* has shown a decline in polyp ingestion rates by ca. 67% at 31°C (67). This matches the observed average drop in C and N assimilation in our experiment (57-66% reduction for the host tissue layers) and also explains the reduced enrichment in the coral tissue structures that degrade and process food material (i.e. the heterotrophic hotspots). The consistent shift in the internal partitioning of nutrients in fed corals for both feeding modes, benefiting symbiont and epidermis, suggests a modulating impact of temperature specific to the gastrodermis as central hub of autotrophic and heterotrophic nutrient assimilation and distribution. This is furthermore supported by the fact that the ~1.5-ratio between epidermal and gastrodermal area was unaltered by temperature in fed corals. Thus, the change in nutrient partitioning was truly related to nutrient allocation and assimilation efficiency and not just due to size changes of the compartments as e.g. observed in *Pocillopora damicornis* at elevated temperature (68). The specific mechanism behind this shift remains unclear. Notably, the symbiont gains more of this increased C and N share under elevated temperature than the epidermis and the NanoSIMS images indicate that this is related to a stronger acquisition of respiratory CO_2_ and organic and inorganic N derived from the food breakdown. Thus, it might be reasonable to assume a stronger diversion of heterotrophic food for catabolic activity in the gastrodermis at elevated temperature, which provides additional energy under elevated temperature for the host on the one hand, but also supports symbiont nutrient fixation and internal recycling. This suggestion of enhanced gastrodermal catabolic activity at elevated temperature should however be treated carefully, since to date it has not been possible to verify this through other proxies for respiratory activity (i.e. metabolic CO_2_ release) that can be measured in a tissue-specific manner. Increased catabolization of heterotrophic C is nevertheless consistent with observations in some corals recovering from bleaching (69, 70). The increased symbiont share of heterotrophic C and N at elevated temperature provides some support for the proposed role of heterotrophic feeding in alleviating symbiont nutrient and CO_2_-limitation, which under bleaching conditions might contribute to preventing sink limitation of photosynthesis and maintaining autotrophic capacity (71).

Contrasting the abundance and distribution of autotrophic and heterotrophic nutrients in the coral symbiosis using NanoSIMS represents a new tissue and cellular-level view of the two main fundamental modes of coral nutrition and shows the tight connection between auto and heterotrophy in a symbiotic coral. This study nevertheless represents a 6 h snapshot of the tissue in a healthy coral known to have a comparably high temperature threshold in its location (39). Thus, the observed role of autotrophy and heterotrophy has to be considered within this context. A number of studies have shown the significant role that heterotrophy plays in the survival and recovery of corals after severe stress events that left their autotrophic capacity impaired (69, 72-76). Thus, while the preservation of symbiont functioning and initial host energy reserves are key elements of coral resistance to thermal stress (77), it is also the complementing interplay of auto and heterotrophy that defines the nutritional state and ability of corals to recover after a severe symbiotic disturbance in a warmer ocean (78).

## Acknowledgements

This work was supported by a Swiss National Science Foundation grant no. CR2312-141048 to AM. The NanoSIMS instrument was acquired with funding from ERC Advanced grant no. 246749 (BIOCARB) and from EPFL to AM. The Red Sea Simulator was funded by an Israel Science Foundation grant to MF. Samuel Donck is acknowledged for help with maintaining the system during the experiments. We thank Dr Emma Gibbin for help with the pyruvate samples. The Electron Microscopy Facility (EMF at University of Lausanne) is acknowledged for access and help with sample preparation; in particular, we thank Dr Jean Daraspe. The manuscript benefited from advice on the histological terminology from Prof Esther Peters.

